# CommonNNClustering—A Python package for generic common-nearest-neighbour clustering

**DOI:** 10.1101/2022.11.28.518169

**Authors:** Jan-Oliver Kapp-Joswig, Bettina G. Keller

## Abstract

Density-based clustering procedures are widely used in a variety of data science applications. Their advantage lies in the capability to find arbitrarily shaped and sized clusters and robustness against outliers. In particular, they proved effective in the analysis of Molecular Dynamics simulations, where they serve to identify relevant, low energetic molecular conformations. As such, they can provide a convenient basis for the construction of kinetic (coreset) Markov-state models. Here we present the opensource Python project CommonNNClustering, which provides an easy-to-use and efficient re-implementation of the commonnearest-neighbour (CommonNN) method. The package provides functionalities for hierarchical clustering and an evaluation of the results. We put our emphasis on a generic API design to keep the implementation flexible and open for customisation.

## Introduction

Density-based clustering procedures—like CommonNN clustering—identify clusters in general as data regions of high sample density separated by sparse, low density regions^1^ and have interesting properties for a wide range of applications. In particular, they are useful in the classification of molecular structures because clusters identified by density-based clustering methods tend to have a natural correspondence to what is understood as a molecular *conformation*: an ensemble of structures with relatively high observation probability associated with the same potential energy minimum or separated by sufficiently small energetic barriers (see figure 1 for an illustrative example).

**Figure 1:**
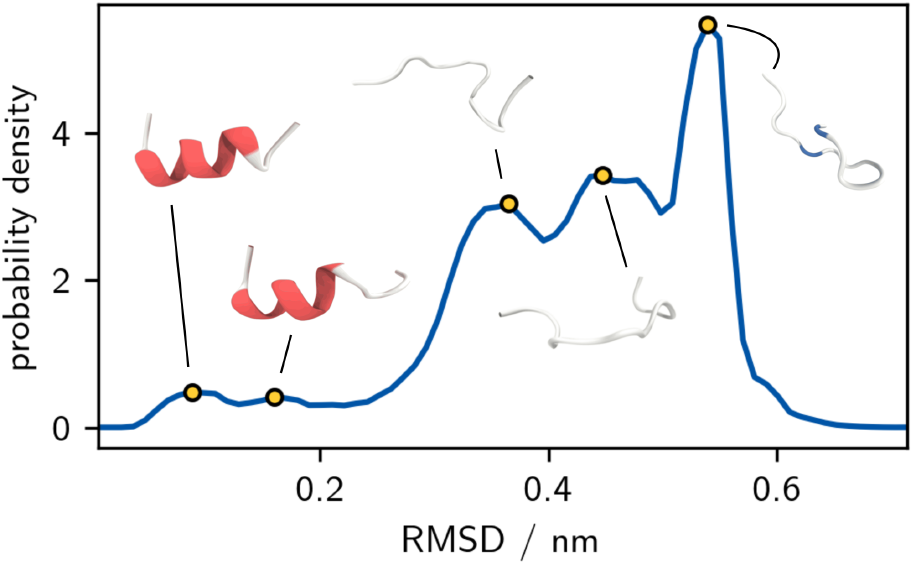
Molecular conformations with low potential energy (high observation probability) identified in a MD simulation of a small helical peptide (PDB ID 6A5J), shown here projected onto the root-mean-square deviation (RMSD) of backbone atom coordinates with respect to the starting structure.

Three points make density-based clustering methods exceptionally suitable in this situation: 1) a conformational cluster is not constrained to a particular shape or layout. Neither is it restricted in its size or extent. CommonNN clustering makes no assumptions in this regard. 2) Not every molecular structure is a good representative for a stable conformation. This means, it is usually beneficial if a clustering can treat individual data points as outliers (noise), which is the case for the CommonNN method. 3) The representations of molecules, i.e. the data space in which they are clustered, can be high dimensional and arbitrarily structured. In general, it is not possible for example to know the correct number of conformational clusters that are to be found beforehand. The clustering should not require any prior knowledge about the data, or should allow easy data exploration. CommonNN offers systematic parameter screening and optimisation, and can decompose a data set hierarchically.

CommonNN clustering has proven to be a viable density-based clustering scheme. It yields intuitively correct clustering results in a wide range of challenging test data cases, as we showed previously in comparative benchmarks.^2,3^ In application, CommonNN clustering has been for example successfully used to characterise the rich conformational ensemble of a foldamer and a tandem WW domain. The latter features two rigid protein domains connected by a flexible linker that sample a huge variety of relative orientations and domain-domain interfaces. ^4^ In other instances, the conformational clustering of small organic molecules has been applied in the context of ligand-protein interactions and pharmacophore modelling.^5,6^ As a discretisation scheme for very well converged core-set Markov models,^7,8^ CommonNN clustering is capable of detecting subtle changes in conformational equilibria and has been useful to explain differences in the membrane permeability of cyclic peptides^9^ and to describe regulative allosteric processes.^10^ Meanwhile, the CommonNN scheme found adaptation in a volume-scaled variant vsCNN,^11^ used in very recent research^12,13^.

In this work, we present a revisited implementation of density-based CommonNN clustering and we provide an accessible Python package to make its use easier and more efficient for a broader audience.

We will quickly summarise the theoretical idea underlying the clustering method and how we approach its algorithmic realisation in section *Theoretical background*. In the sections *Basic usage, Advanced usage*, and *Practical advice* we describe the usage of the package and some of the program design decisions. Section *Benchmark* provides a small benchmark of the new implementation.

### Theoretical background

Consider a data set of points that should be clustered as a set of samples from an underlying probability density *p* : Ω → ℝ_≥0_ with respect to a *d*-dimensional feature space Ω ⊂ ℝ^*d*^. In the case of molecular data, Ω is a configurational space of structural features and *p* is a corresponding configurational Boltzmann density. Density-based CommonNN clustering can be formulated resting on the idea of applying a density threshold *λ* to *p* that separates Ω into regions of high and low density like shown in figure 2 for 1D. Clusters are the resulting isolated, continuous regions of high density while everything below *λ* is noise. The set of possible clustering results is systematically gathered through variation of *λ*.

**Figure 2:**
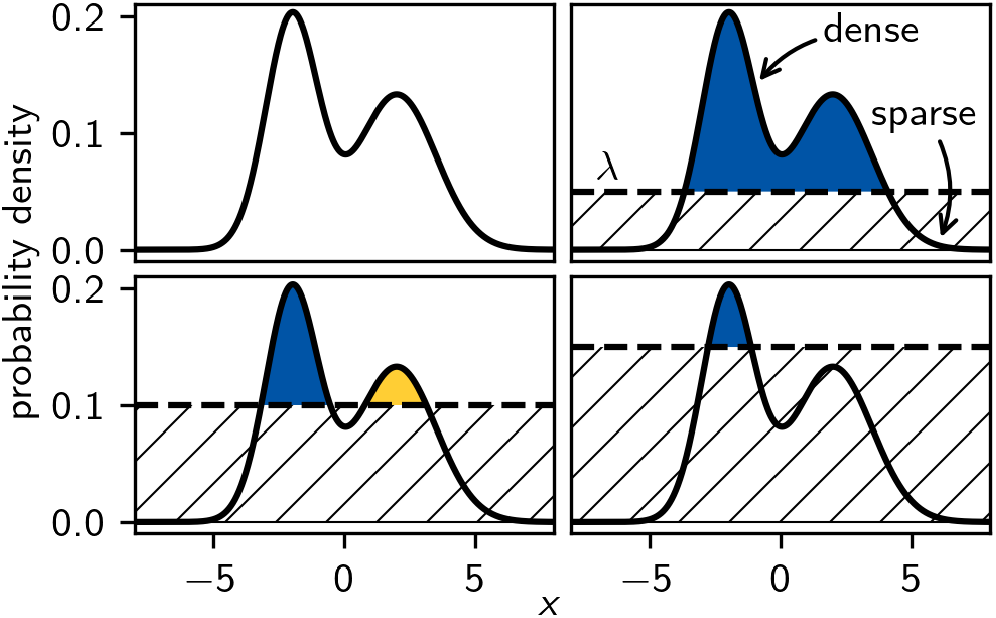
Example probability distribution in 1D with two maxima. Applying different density-thresholds *λ* splits the distribution into isolated high-density regions (clusters) separated by low density (noise). Starting with a low threshold of *λ* = 0, all data points are assigned to the same single cluster. Higher thresholds lead to low density points to fall into the noise region, existing clusters to shrink, and eventually to splits if a local minimum is exceeded.

CommonNN clustering employs a local estimate of the density, based on discrete point samples: the proxy for the density is the number of data points within the (fixed radius) neighbourhood intersection of two points, i.e. the number of common nearest neighbours. A threshold (compare *λ*) on this density estimate can be given as a minimum number of *n*_c_ common neighbours. Additionally, a point pair is *connected* through the density estimated in terms of a *similarity* measure. Two points for which the density-threshold is exceeded are identified to be part of the same cluster (see figure 3). The two points are in this case said to “fulfil the density-criterion” or “pass the similarity check”. The density estimate usually requires the definition of a metric on the feature space that allows the calculation of pairwise distances and the determination of neighbourhoods.

**Figure 3:**
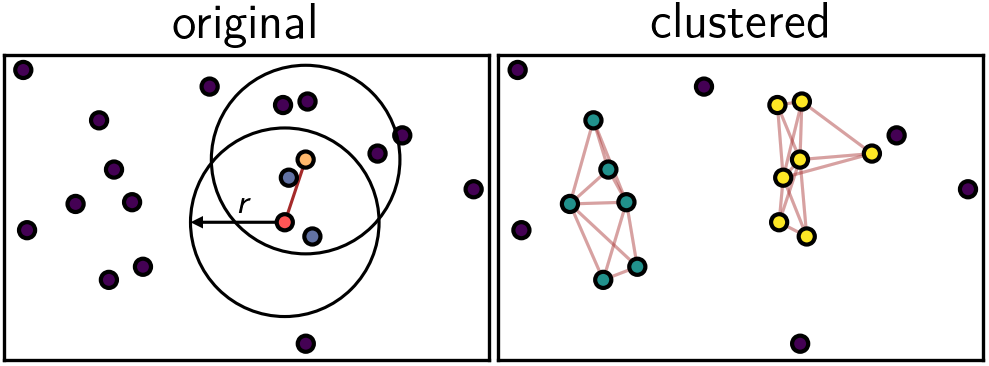
Illustration of the density-criterion in the CommonNN scheme for random points in 2D. The red and the orange data point (left) share two of their neighbours with respect to the search radius *r* (blue points). For a set value of *n*_c_ ≤ 2, the two points are considered part of the same dense region (thick red edge between them). A sub-network of points connected in this way (right) constitutes a cluster (yellow and green points).

It is convenient to think of a data set that should be clustered as a graph *G*(*V, E*) in which each point is represented by a node (vertex) *v*_*i*_. Edges *e*_*ij*_ indicate pairwise relationships between points. If the edges correspond to whether two points *v*_*i*_ and *v*_*j*_ fulfil the density-criterion, the connected components— sub-networks of nodes that are disjoint from the rest—of the graph are the clusters that we want to find. Hence, the main task of the clustering can be solved by leveraging well established graph traversal algorithms, for example a breadth-first-search approach (see figure 1 in the SI).

### Basic usage

The CommonNNClustering package requires Python ≥ 3.6. It can be installed from PyPi (`pip install commonnn-clustering`) or from the development repository on GitHub (https://github.com/bkellerlab/CommonNNClustering). The installation requires Cython, which is used to implement core functionalities efficiently. At runtime, NumPy is mandatory as well. Optionally, Matplotlib, Networkx, Pandas, scikit-learn, and scipy are leveraged for additional functionality. Documentation is available under https://bkellerlab.github.io/CommonNNClustering. An alternative implementation is furthermore available within scikit-learn-extra (`github.com/ scikit-learn-contrib/scikit-learn-extra`).

Getting started with the clustering of any data set using the CommonNNClustering package is easy and should feel familiar to the use of similar available object-oriented Python APIs like that used by scikit-learn.^14^ The code snippet in figure 4 illustrates the four essential steps of a clustering: 1) how to import the main `cluster` module (line 1), 2) prepare a clustering object as an instance of the `Clustering` class from the data (line 5), 3) trigger the clustering (`fit`) itself with specified parameters (line 6), and 4) access the resulting cluster label assignments for further analysis (line 7). As a general design principle, we settled on a data oriented approach for the whole clustering procedure, which means that a created clustering object will be always associated with exactly one data set of some form. This data set can be clustered based on different combinations of cluster parameters, re-using the same clustering object.

**Figure 4:**
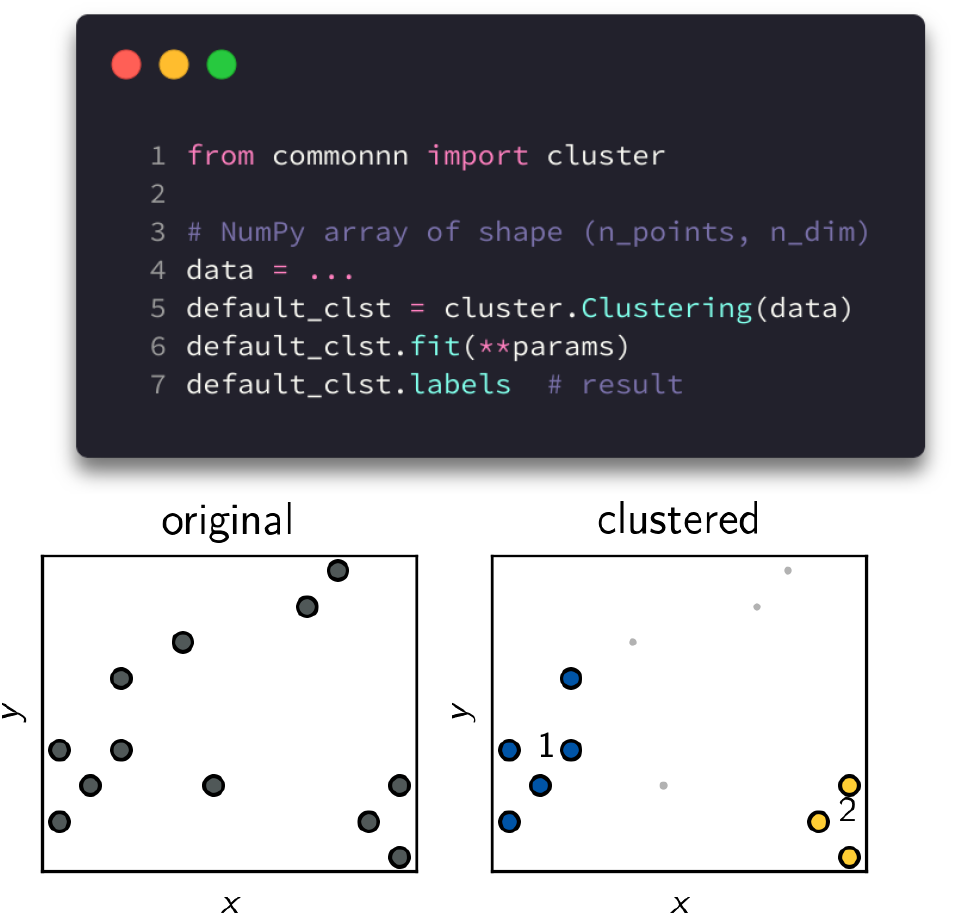
Default cluster object creation and data point clustering. The scatter plots below are created using the convenience method `default_clst.evaluate()`.

The example assumes the presumably most frequent use-case of having the data presented as a NumPy array of shape (#points, #dimensions) or something equivalent for that matter, the data contains information on feature space coordinates for each sample point. The program can use these for the calculation of distances and effectively neighbourhoods to perform the clustering. This is, however, not the only possible scenario. In general, input data can be fed into a clustering in one of three fundamentally different formats: 1) point coordinates, 2) pairwise inter-point distances, or 3) fixed radius neighbourhoods. Each of these basic types of information can eventually be materialised in a multitude of different data structures. In particular, it is allowed and encouraged to leverage other specialised programs to take over the distance or neighbourhood calculation, e.g. with *kd*-trees as provided by scikitlearn.^14^

### Advanced usage

We designed the CommonNNClustering code so that it is flexible with regard to different input formats and variations in other constraints, like for example the used distance metric. This allows the user to customise the program to individual needs, also beyond the application to standard MD data. This section and Figure 5 give a short overview of the design architecture of the package. The section may be skipped if the default behaviour described in the last section is sufficient.

**Figure 5:**
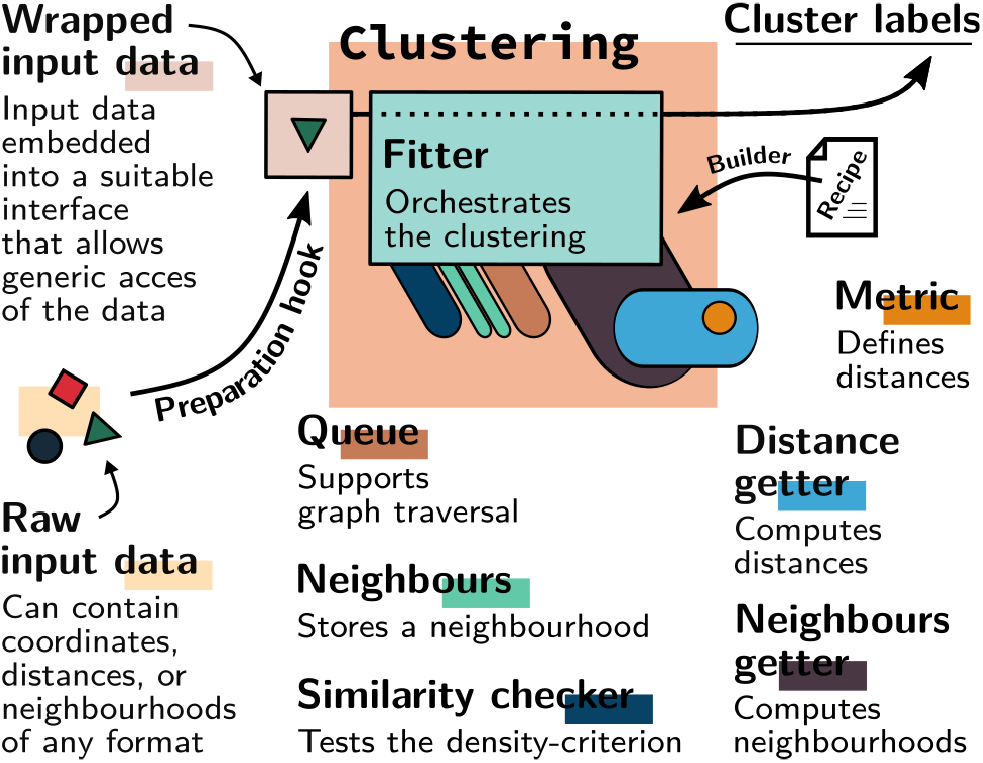
Aggregation of a clustering object from generic types representing exchangeable clustering components. A so called *fitter* implements the clustering procedure, making use of generic interfaces.

The central idea is the following: when the clustering procedure is executed, it has to loop over input data points and query their neighbourhoods. Instead of accessing whatever input data structure is presented directly, the raw input data is wrapped within one of several *input data* objects that all can be worked with through a common generic interface, i.e. which are of a certain defined type. The task of acquiring neighbourhood information is delegated to one of many possible *neighbours getter* objects. In this way, the clustering that is itself implemented in a *fitter* object does not have to be concerned with how needed information is stored in and retrieved from the input data. In the same way, other important components as the testing of the density-criterion (*similarity checker* type), intermediate storage of retrieved neighbourhoods (*neighbours* type), or the metric (*metric* type) used for distance calculations (*distance getter* type) are represented by exchangeable objects adhering to generic interfaces. A clustering object as initialised in the code example in figure 4 aggregates all the objects needed for a clustering and is assembled in the background according to a *recipe*.

Please refer to the documentation for details on how the different interfaces are defined exactly, which generic types are available already, and how custom types and recipes can be defined and invoked.

### Practical advice

The outcome of a CommonNN-clustering depends on the two cluster parameters *r* (neighbour-search radius) and *n*_c_ (CommonNN density-cutoff). For higher values of *n*_c_ at a given *r*, two points are assigned to the same cluster only if the respective pairwise density estimate is high enough. Which values are eventually to be chosen for *r* and *n*_c_ depends strongly on the (subjectively) expected clustering result and on the nature of the data set (its distribution and sampling), and is in general not possible to decide *a priori*.

In selecting a suitable radius *r*, the aim is to choose a value that allows for a sufficiently local density estimate. In this sense, *r* functions as a “resolution” for the clustering. With a low resolution (a large radius *r*), local differences in the point density can not be detected, and no splitting of the data into clusters is achieved. If *r* is very small, fluctuations in the local point density that may originate from insufficient sampling, can lead to meaningless splittings of points into undesired clusters. Furthermore, with small values of *r*, the sensitivity of the density threshold towards *n*_c_ is increased. As a heuristic for a good first guess on a neighbour search radius *r*, which allows an appropriately local density estimate, it has proven useful to take the distance value at which the distribution of pairwise inter-point distances has its first maximum.

To determine *n*_c_, we use the strategy of setting *r* according to said heuristic and screening *n*_c_ from small to large values, to obtain clusterings at increasingly high density thresholds.^2^ This allows to systematically select the favoured clustering result. For comparison, *r* may be adjusted slightly in both directions— in particular for high sampling rates, the resolution can usually be increased. Figure 6 illustrates this strategy with three representative data sets. When settling on a specific clustering result, it is advisable to look for a parameter range in which the clustering is qualitatively stable, meaning where only the size of individual clusters varies (shrinks for increased *n*_c_) but not their number. The general observation is that while the density-criterion goes up, the number of clusters increases as more and more splittings occur. The number will eventually go down again as more and more low density clusters fall below the density threshold and vanish into noise.

**Figure 6:**
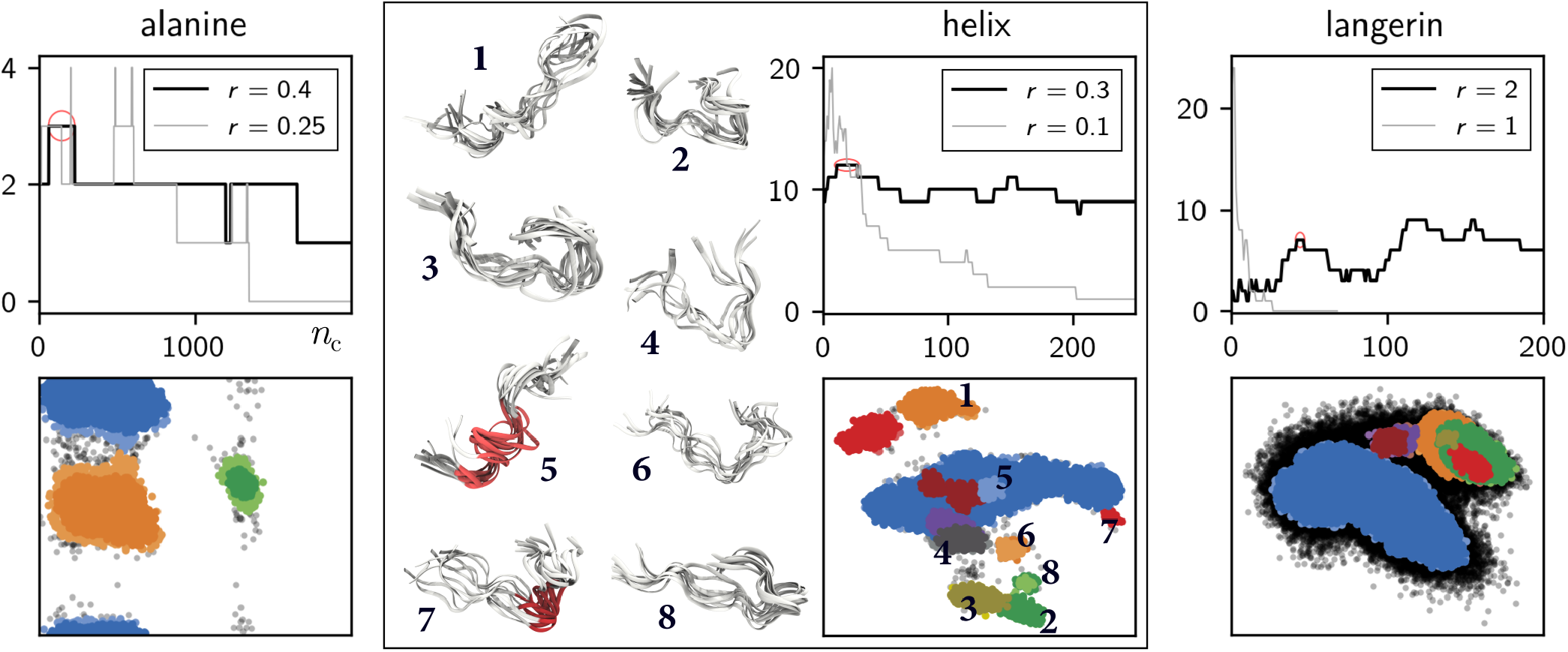
1D cluster parameter scans: number of clusters vs. *n*_c_ for fixed radii *r* (proposed initial guess and much smaller). The most promising regions (stable, high cluster number) are marked with red. Clustering examples for parameters in the highlighted ranges below. Data points are coloured by cluster label in two shades of the same colour for the lowest and highest *n*_c_ value, respectively. See SI for methods.

CommonNN clustering is intrinsically hierarchical (compare figure 2). The variation of the density-criterion essentially leads to a hierarchy of clusterings, and individual clusters may be extracted at different levels of this hierarchy. Specifically, one may want to further split higher density clusters without loosing low density clusters into noise. We support currently two ways of doing this around threshold-based clustering: a *manual* and a *semi-automatic* approach.

Full user-control on each hierarchy level is offered by the manual approach as illustrated in SI figure 2 and figure 7. The idea is to follow the strategy of increasing *n*_c_ with fixed *r* to a point where the number of isolated clusters is locally maximised without too many low density clusters being lost into noise. Than this clustering result is isolated, i.e. frozen or saved, and the clustering is continued only on a subset of child clusters. Finally, the resulting hierarchy can be reeled—wrapped up—back into a single partitioning.

**Figure 7:**
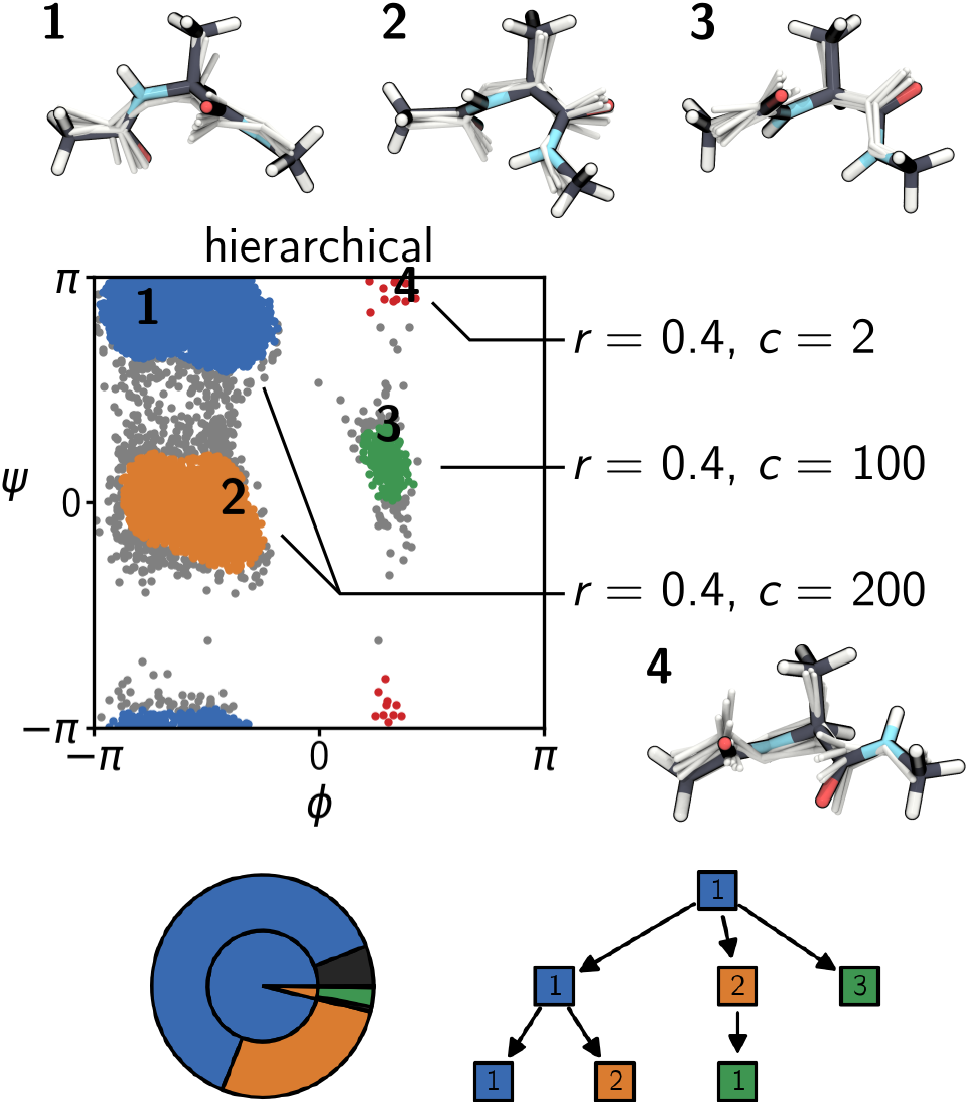
Manual hierarchical clustering of the *alanine* data set. Upper: Data points coloured by cluster label assignment after 2-step clustering (reclustering of cluster 1 and 2 obtained in a first step). Lower: A hierarchy of clustering objects can be visualised as pie-chart showing the hierarchy levels going outward from high (root) to low (children) where the size of the pieces represents the amount of data points in a certain cluster. Alternatively, a Sugiyama tree-diagram can be drawn showing the splittings at individual hierarchy levels from top to bottom.

The second approach is based on the idea to semi-automatically build the hierarchy of clusterings at certain levels by specifying a list of parameter combinations. This necessitates that in a second step, the resulting hierarchy needs to be screened according to some criteria for which child clusters should be kept as the final result. SI figure 3 and figure 8 illustrate this approach employing a hierarchy screen followed by a trimming to avoid that clusters shrink (and eventually vanish completely) if they do not split further. Again, please refer to the documentation for more details on hierarchical clustering and different trimming approaches.

**Figure 8:**
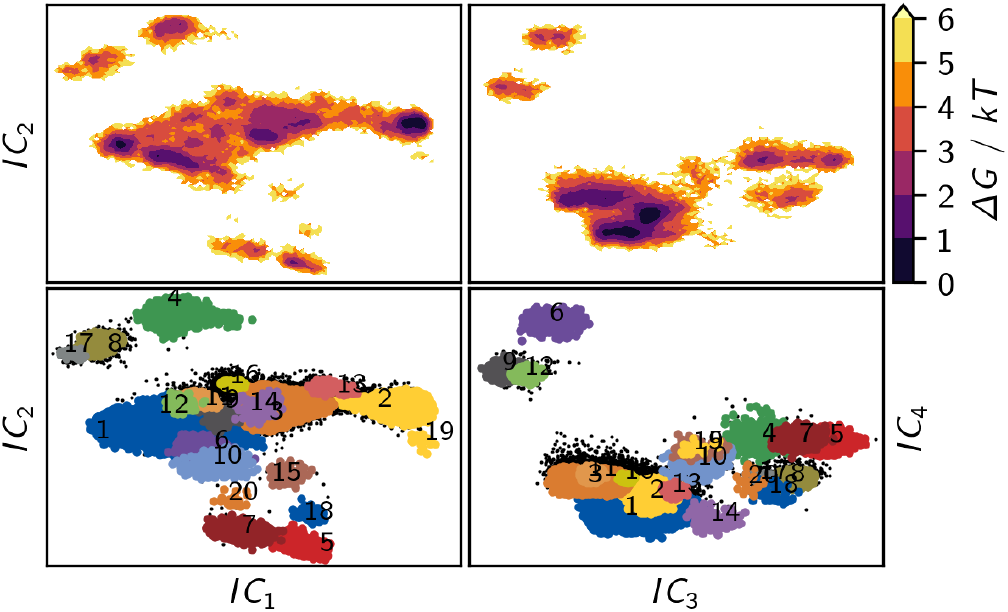
Semi-automatic hierarchical clustering on the *helix* data set. Upper: Pseudo free energy surface of the distribution. Lower: Data points coloured by cluster labels (*r* = 0.3, *n*_c_ ∈ [0, 600]) after trimming.

### Benchmark

If large data sets are clustered or many parameter combinations are used, it becomes important that the clustering is relatively fast. Benchmarking the computational efficiency of a clustering is difficult because it does not only depend on the implementation itself but also on the type and structure of the data set. The timings depend non-trivially on the cluster parameters, which influence how often neighbourhoods need to be retrieved and checked, and how quickly this can be done. CommonNN clustering can also be realised in several different ways (using different combinations of generic types). Besides pure computational efficiency, memory concerns may also be of importance.

We show the clustering performance with collected timings under varying constraints using qualitatively the same data set but an increasing number of points. Empirical scaling was determined by fitting the measured execution times *t* versus problem size *n* to the power function *t* ≈ *an*^*b*^ where *a* is a proportionality constant and *b* is the growth factor of interest. The presented timings have been measured on a Debian 10 operating system, equipped with an Intel(R) Xeon(R) CPU E5-2690 v3 @ 2.60GHz and 164 GB RAM. Note, that all implementations are serial at the moment. Timings are reported as the best out of ten repetitions. Memory usage is reported as allocated resident memory and the lowest out of ten repetitions.

Figure 9 shows a benchmark on uniformly distributed points with fixed cluster parameters. The CommonNN-cutoff is set to *n*_c_ = 0, which means that the similarity criterion check is essentially skipped and the timings reflect only the general breadth-first search clustering procedure, including the construction of the intermediate neighbour lists. This allows us to derive a cluster parameter independent performance baseline. We compare different input data formats and the corresponding default recipes (see the documentation for details on what these recipes entail). In all cases, we observe an empirically quadratic scaling of the computation time with respect to the number of points in the data set. By using pre-computed distances or neighbourhoods as the input, one can save a substantial amount of computing time. Neighbourhoods sorted by point indices give absolutely the best performance: 256,000 data points can be clustered on the order of seconds.

**Figure 9:**
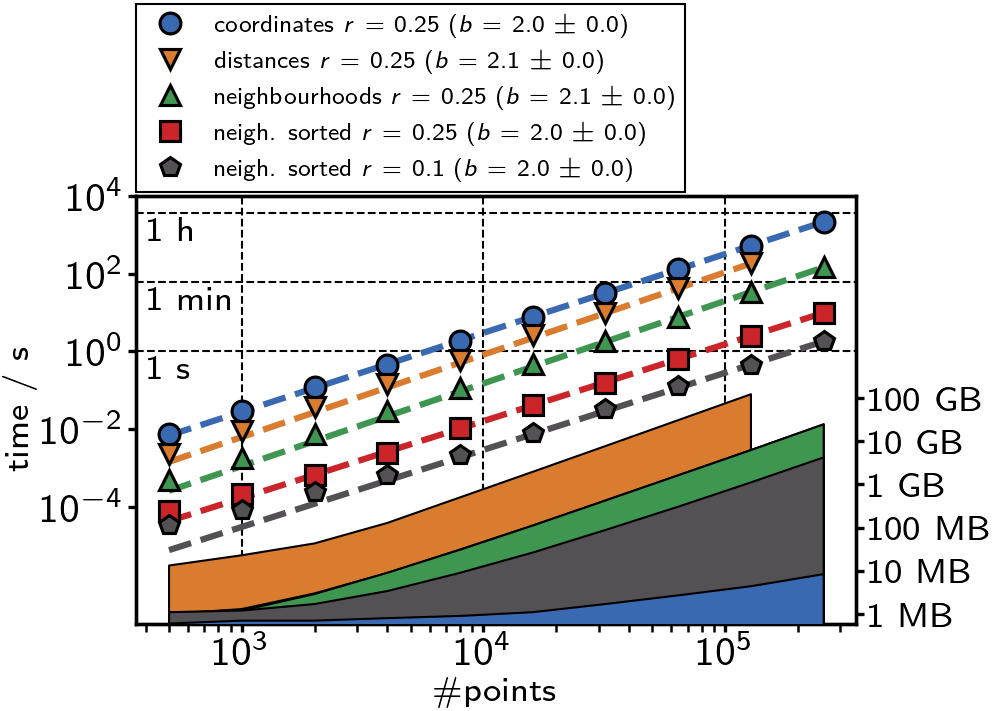
CommonNN performance for a data set with an increasing number of uniformly distributed points in 2D (see SI) using different default recipes. Execution time measurements (dashed lines) include the clustering only, disregarding data preparation. The similarity cutoff was set to *n*_c_ = 0 in all runs. Memory demand (filled areas) covers the complete Clustering objects, including input data, during the clustering.

Pre-computed information occupies a certain amount of memory. In particular the storage of a square distance matrix can become prohibitively expensive. Note that for 256,000 data points, the available memory was exceeded in our case. Pre-computed neighbourhoods require much less memory. Furthermore, we see that it is beneficial to choose a rather small radius *r*. This will keep the number of neighbours per point small, which leads to faster filling of the neighbours containers and also to a lower memory demand, especially if neighbourhoods are pre-calculated.

To evaluate the cost of input data preparation, figure 10 shows benchmarks including the time needed to pre-calculate distances or neighbourhoods. The advantage of using precalculated distances is essentially nullified if the preparation time has to be considered for the overall clustering performance. Clustering from pre-computed neighbourhoods clearly outperforms starting from point coordinates even if the preparation time is considered. Here, putting in the effort of sorting the neighbourhoods still offers a little edge. The precomputation is amortised already in the first run. For these benchmarks, the cluster parameter *n*_c_ was fixed at a non-zero value so that similarity checks are included. The radius *r* was scaled down progressively, which can be justified for large (well sampled) data sets. This has the effect of producing sub-quadratic scaling.

**Figure 10:**
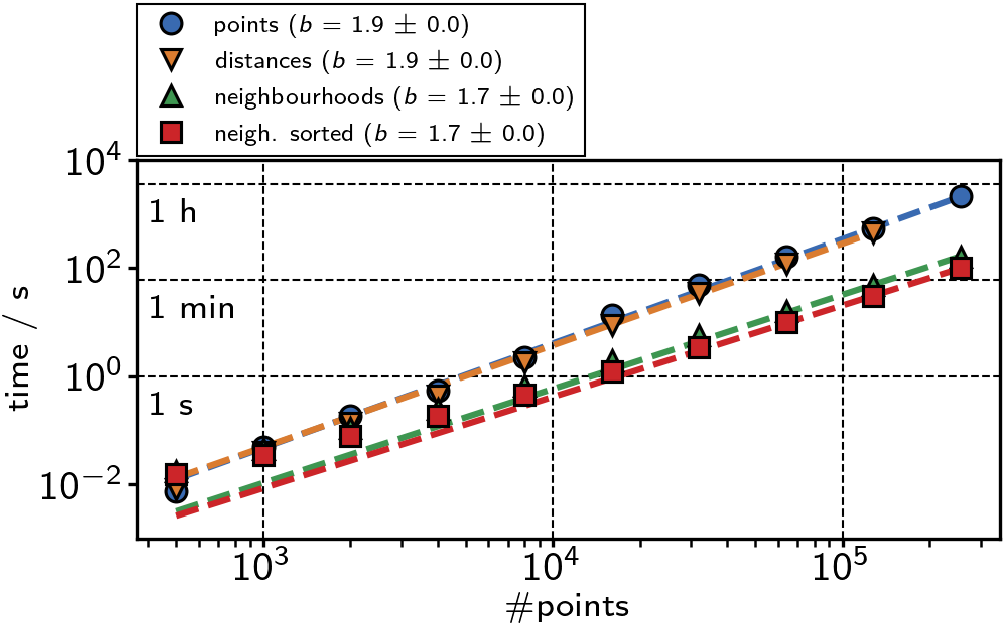
Full CommonNN clustering benchmarks including input data preparation time, based on the *varied* data set (see SI). The similarity cutoff was set to *n*_c_ = 50 while the neighbour search radius was set to *r* = 0.2 initially and scaled down by a factor of 0.9 each time the number of points was increased.

In summary, it is generally not recommended to use a pre-computed distance matrix as input source. Pre-computed neighbourhoods are more beneficial in terms of execution time and memory demand, although they have to be recomputed for changing radii *r*. If memory is very limited, plain point coordinates may be the best option.

## Conclusion

We demonstrated the `commonnn` Python package that provides a convenient user interface to threshold-based, density-based CommonNN clustering. The presented revised implementation rigorously improves our previous one in terms of clustering performance, usability and flexibility. Its generic design allows the application of the procedure in a wide range of situations. The package is open to be extended with specialised types to cover additional use-cases— for example other forms of input data. Future work will be dedicated to broaden the array of available types and on incorporating computational parallelisation schemes into their design. Furthermore, automatic hierarchical clustering without explicit use of a threshold is explored. Currently, we also refine the application of the procedure to dynamic pharmacophore data to cluster protein-ligand complex binding poses.

## Acknowledgement

Funded by the Deutsche Forschungsgemeinschaft (DFG, German Research Foundation) under Germany’s Excellence Strategy – EXC 2008/1 – 390540038 and through grant GRK2473 ”BioactivePeptides” - project number 392923329.

