## supporting information for "CommonNNClustering—A Python package for generic common-nearest-neighbour clustering"

### Breadth-first-search graph traversal

```
Init cluster label (to 1)

for node in graph:
  If node is visited:
    continue
  # New component
  Pick source node
  Mark node visited
  Push node to queue

  # Start breadth-first-search
  while queue is not empty:
    Pop node off of queue
    for node in connected_nodes:
      If node is visited:
        continue
      Assign label to node
      Mark node visited
      Push node to queue

  Increment cluster label
```

Figure 1: Pseudo code for a breadth-first-search graph traversal to identify connected components (clusters) in a data graph. The CommonNN density-criterion determines if two nodes are connected.

### Hierarchical clustering

```
hier_clst.fit(**params)
hier_clst.isolate()

recluster_list = [(1, params_a),
                  (2, params_b)]
for subcl, params in recluster_list:
  hier_clst.children[subcl].fit(
    **params)

hier_clst.reel()
```

Figure 2: Manual hierarchical clustering involves the isolation of clustering results and a re-clustering of child clusters.

```
r = 0.3
c = [1, 50, 100, 200, 500]

hier_clst.fit_hierarchical(
  r, c, member_cutoff=20
)
hier_clst.trim()
hier_clst.reel()
```

Figure 3: Semi-automatic hierarchical clustering is based on the specification of a list of cluster parameters and selection of child clusters according to user-defined criteria.

### Molecular dynamics data sets

The example data sets referred to in the main article (section *Practical advice*, figure 6) have the following sources:

- ***alanine*** (10020 points, 2 dimensions): 4 µs of Ac-A-NHMe (20 replica á 200 ns) in TIP3P water using the AMBER99SB-ILDN force field were produced for *"Dynamic Properties of Force Fields"*, F. Vitalini, A. S. J. S. Mey, F. Noé, and B. G. Keller *J. Chem. Phys.* **2015**, *142*, 084101 (`https://doi.org/10.1063/1.4909549`). The raw simulation data have been projected onto backbone dihedral angles as described therein.

- ***helix*** (37500 points, 4 dimensions): 3.75 µs of a small peptide structure with PDB ID 6A5J (5 replica á 750 ns) in TIP3P water were produced using the AMBER14SB force field in OpenMM 7. The simulations were driven by a Langevin integrator ($T = 300\,\mathrm{K}$, $\gamma = 1\,\mathrm{ps}^{-1}$, $\Delta t = 2\,\mathrm{fs}$), using constraints on bonds involving hydrogen atoms, PME electrostatics, and a non-bonded interaction cutoff of 1 nm. The system has been solvated, neutralised, minimised, and equilibrated in the $NVT$ ensemble for 100 ps prior to production. We thank Puneeth Kouloorkar for producing the raw simulation data during an internship in the group. Subsequently, time-lagged independent component analysis on backbone, $\chi_1$, and $\chi_2$ torsion angles ($\tau = 5\,\mathrm{ns}$) was employed to project the data.

- ***langerin*** (103853 points, 6 dimensions): 27 µs of langerin (117 replica á 150 to 1000 ns) in TIP3P water using the AMBER99SB-ILDN force field were produced for *"The molecular basis for the pH-dependent calcium affinity of the pattern recognition receptor langerin"*, J.-O. Joswig, J. Anders, H. Zhang, C. Rademacher, and B. G. Keller *J. Biol. Chem.* **2021**, *296*, 100718 (`https://doi.org/10.1016/j.jbc.2021.100718`). The raw simulation data have been projected using a principle component analysis on $C_\alpha$-atoms as described therein.

### Benchmark data sets

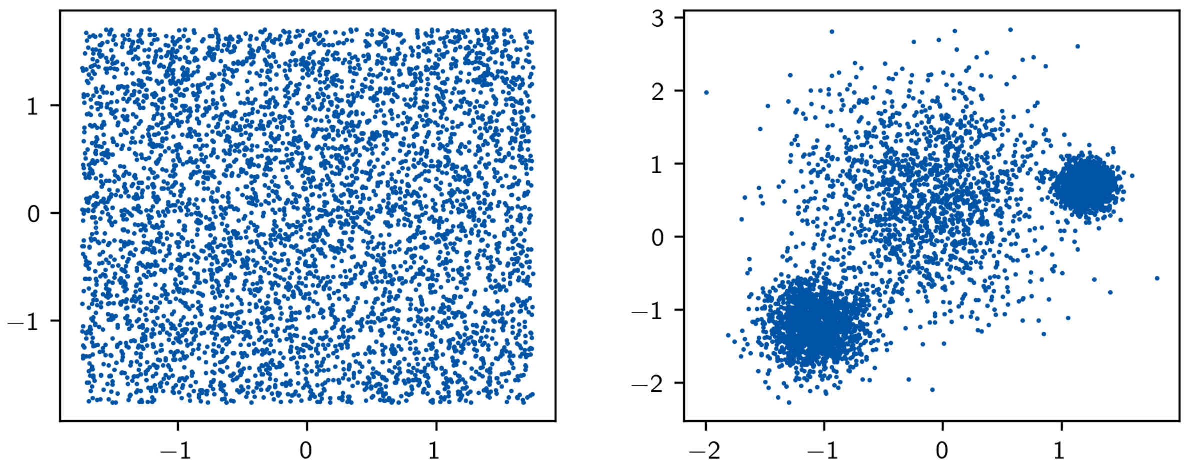


Figure 4: Left: 5000 uniformly distributed random data points as used for figure 11. Right: 5000 *varied* data points generated with `sklearn.datasets.make_blobs` as used for figure 12.
